# A Dose-Response Model and D_10_-Value for *Mycobacterium tuberculosis* Exposed to Dosimetrically Verified Ionizing Radiation

**DOI:** 10.1101/2021.03.02.433681

**Authors:** Jackson V Watkins, Justin Bell, Phillip Knabenbauer, Alexander Brandl, Karen M Dobos

**Author notes:** Contributed equally to this work (Jackson V Watkins, Justin Bell).

## Abstract

Techniques for pathogen inactivation have been employed by laboratories to help ease the financial, physical, and health strains associated with (A)BSL-3 work. Exposure to radiation is the most common and useful of these methods to inactivate pathogens grown in large-scale culture. While robust protocols exist for radiation exposure techniques, there are variances in methods used to determine the radiation dose and dose rate required to inactivate pathogens. Furthermore, previous studies often do not include radiation dosimetry verification or address corresponding dosimetry uncertainties for dose response-assays. Accordingly, this study was conducted with the purpose of completing a dosimetry assessment of the radiation field within the sample chamber of a sealed source irradiator, to subsequently determine the radiation dose required to inactivate pathogenic cultures. Physical dosimetry techniques (Fricke dosimetry, ion chamber measurements, and measurements with thermoluminescent dosimeters) were used to measure dose rate and rate variances within the sample chamber. By comparing the variances between the dosimetry methodologies and measurements, an estimated dose rate within the sample chamber was determined. The results of the dosimetry evaluation were used to determine the radiation dose samples of *Mycobacterium tuberculosis* received, to accurately associate biological markers of inactivation to specific doses of ionizing radiation. A D_10_ value and dose-response curve were developed to describe the inactivation of *Mtb* from increasing doses of ionizing radiation. The D_10_ value is experimentally relevant for comparative analysis and potentially provides a biological baseline for inactivation verification. This methodology can also easily be translated to other pathogen models.

**Importance:** This work set out to give us a better understanding of how much radiation is required to inactivate *Mycobacterium tuberculosis*, the bacteria that causes tuberculosis disease. Radiation dose from a source is not something that can just be inputted, it must be calculated, so we also determined the approximate dose from the source to address ambiguities that had previously existed while inactivating microbes. We were able to generate an accurate description of inactivation of *Mycobacterium tuberculosis* by correlating it with a value representing 90% death of the treated cells. We also unexpectedly discovered that very low levels of radiation increase certain activity within the cell. This is important because it allows us to better understand how radiation kills *Mycobacterium tuberculosis*, and gives us a value to compare to other organisms. It also offers other researchers a method to use under their own specific conditions.

## Introduction

### Inactivated Pathogens

Human risk group-3 (HRG-3) pathogens maintain clinical and ecological interest because of their ability to infect hosts and manifest diseases with high mortality rates. While there remains a strong desire to study these pathogens for the advancement of human health, proper protection of those who work with these organisms is challenged by financial and mechanical barriers^1^. For researchers to mitigate risk of laboratory acquired infections (LAI), respiratory pathogen work requires high-containment settings such as biosafety level-3 (BSL-3) laboratories for the safe growth and manipulation of such organisms^2^. While these laboratories are most ideal for protection of both researchers and their communities, the aforementioned financial burdens make them nearly infeasible in some parts of the world^3^. It too must be acknowledged that any facility conducting work with deadly and contagious pathogens, regardless of financial means, maintains some risk of spill-over from the laboratory setting into the communities where the research is situated^4,5^. Each of these barriers to high-risk pathogen research are amplified in regions which are resource-poor, wherein those same regions find themselves most susceptible to outbreaks^1,3,6^. And though these barriers are certainly felt strongly in resource-poor communities, they are present even in communities with ample means to finance research, as facilities require maintenance and extensive training, while having a slight but non-zero risk of infection to researchers and subsequently, their communities^7^.

### Inactivation Techniques

A wide variety of inactivation techniques have been developed for the purpose of rendering pathogens non-viable, while also leaving cell constituents, media, or suspension of interest intact^48^. Such techniques for sterilization are useful in food and water-system safety^8-10^, purification of transfusion and plasma-derived products^11-13^, and laboratory applications such as diagnostics and basic science research^14,15^. In the context of the latter, inactivation techniques are beneficial for the continuing safety of those conducting research with biological hazards. Our interest in the application of inactivation techniques stems from a desire to safely work with pathogens and their constituents outside of BSL-3 containment, by applying methods that inactivate the pathogens while limiting the disruption to structural and enzymatic characteristics. Dependent on the research objective, each technique for inactivation can causes positive and negative effects, and a one-size-fits-all approach has not been fully developed. Accordingly, the adopted inactivation method should ideally provide adequate inactivation of the pathogen, while limiting effects to the underlying biological being studied.

Heat inactivation of pathogens is most commonly used for food and water safety purposes but is also utilized in laboratory settings^8-10^. High temperature treatment, especially beyond 95°C, is profoundly effective at inactivation of not only pathogens, but resilient spores formed by highly resilient organisms^8^. Lower temperature treatments have more variability in efficacy of inactivation, but adequate temperatures to inactivate are described in the literature for many common laboratory-strain pathogens^9,10^. As a result, information regarding D_10_ values (commonly referred to as D-values in the literature; the treatment parameters required to reduce bacterial or viral load by 90%) is accessible for a variety of organisms. The comparison of these values has led to robust relative measurements of thermal resistance across microorganisms. However, downstream application for heat-treated pathogens is minimal, as hyperthermic exposure causes widespread damage to proteins, lipids, and many cellular components^8^. While heat inactivation is convenient for DNA preparations due to high temperature requirements for DNA denaturation, it is seldomly used in other contexts where hyperthermic exposure may disturb protein secondary and tertiary structure, cellular component integrity, or normal lipid and enzyme function. This technique, despite its incredible utility in sterilizing instruments or reducing microbial load in food and water products, maintains limited practical use in the laboratory setting.

Lipid solvents have also been well described for bacterial and viral inactivation^12,13^. Solvents like chloroform, hexane and ethyl-ether are useful for isolation of cellular DNA, lipids, and membrane proteins. In low concentrations, chloroform has even been used to reduce pathogenic load in plasma products^13^, though due to the toxicity and hazards associated with many solvents, they can’t be realistically upscaled to large concentrations of cells without generating proportionally large amounts of hazardous waste. Besides the generation of toxic byproduct, another downside is that these methods are only useful for enveloped viruses, or organisms with lipid membranes, made soluble by organic solvents. Lipid solvents are so ineffective at inactivating non-enveloped viruses in fact, that treatment with chloroform and subsequent plaque assays have been used by virologists to classify newly discovered viruses as either enveloped, or non-enveloped^13^.

Photochemical methods have also gained popularity as methods for inactivation of pathogens^16-18^. Photochemistry involves the use of a photosensitizing agent, including but not limited in the literature to riboflavin^16,18^, radachlorin^19^, or methylene blue^19^. The basic principles of photochemistry are similar to those of exposure to ionizing radiation, however ultraviolet (UV) radiation is non-ionizing. The agent must absorb light at a designated wavelength to enter its triplet energy state, which is then transferred to a target at the return to ground state^19^. The target could be oxygen, generating reactive oxygen species, or cellular components such as DNA or proteins, disrupting the normal biological function of the cell and leading to inactivation of the microorganism. In either case, the photochemistry leads to inactivation of pathogens. Mirasol Pathogen Reduction Technology (MPRT) is a common method for mediation of such reactions, and has previously been demonstrated to completely inactivate pathogens in human blood^16-18^, currently making it a useful tool in hospitals for quality control (QC) during blood transfusions^16,17^. Additional studies have also demonstrated complete inactivation and extensive DNA damage in 8 log cultures of *Escherichia coli (E. coli)*^20^. The process of inactivation involves a photosensitizing reaction between UV radiation and riboflavin, which causes 8-oxo-guanosine modifications at guanosine DNA nucleosides. Irreversible double-strand breaks in cellular DNA are characteristic biomarkers of this type of damage^16,20^. UV radiation causes thymine dimers and damages cellular DNA even without the addition of a photosensitizing agent. The benefits of a technology such as this are apparent in the specificity of damage and complete lack of toxic byproduct generation. These attributes are what make the MPRT so useful in hospital settings, as most of the blood product remains intact, while also limiting the toxic by- product which would severely impact patient safety. However, preliminary work in our lab has demonstrated that while a significant reduction in replicating *M. smegmatis* cells could be consistently achieved over multiple replicates, surviving populations of cells remained, even after high doses of UV radiation^21^. So, while the technology has proven to be useful for inactivation of lower pathogen loads in hospital settings, its utility in the laboratory settings is yet to be determined. This is particularly apparent when dealing with pathogen inactivation from large-scale culture.

Commonly, in each of the methods described, there are attributes which prevent broad utility, especially for inactivation of organisms from large-scale culture. Consequently, the financial and mechanical barriers to pathogen research persist.

### Exposure to Ionizing Radiation

Exposure to ionizing radiation has been shown to inactivate pathogenic organisms for safe use outside of a BSL-3 laboratory setting. Specifically, exposures to ionizing radiation inactivate organisms^4,22-25^ through either indirect mechanisms of oxidative stress from the radiolysis of water surrounding the DNA, or by damage from direct interactions between cellular DNA and incident radiation^24,25^. Dependent on the radiation dose cells may remain mostly in tact due to the specificity of the damage, maintaining much of their structural and component integrity, as well as metabolic function^26^. This is particularly useful for shipment of product and applications such as subcellular fractionation, protein and lipoglycan purification, and associated proteomics and metabolomics analysis. The process of exposing HRG-3 pathogens to specific doses of ionizing radiation has allowed for significant advancements in research. These advancements include but are not limited to steps forward in diagnostic techniques^27,28^, vaccine production^29-31^, and basic research immunology^31,^ which all have implications for human health in the face of epidemics and spread of HRG-3 pathogens.

It should, however, be noted that accidental release of inadequately inactivated organisms has been documented as recently as the 2015 Dugway incident^32-34^. During this particular event, samples containing live anthrax spores were incompletely inactivated and shipped from a contained environment, posing a significant risk to both researchers and the public^32^. The event lead to the temporary closure of nine biodefense laboratories, and 41 potential exposures to live samples^32,33^. Such incidents, especially in such recent history, demonstrate both a need for strict protocols when dealing with deadly organisms, as well as require a continued understanding of the methods required to validate and ensure devices used for inactivation of harmful pathogens are functioning properly.

To address these recent concerns, this study aimed to improve the understanding of current inactivation protocols. The primary goal of this research was to develop a D_10_-value for *Mycobacterium tuberculosis (Mtb)* specific to exposures of ionizing radiation. Prior to this study, limited data existed that correlated specific viability markers for *Mtb* or *M. smegmatis* to specific doses of radiation. To support the primary goal of this research, a multitude nuclear instrumentation and radiation detection techniques were employed to characterize the radiation field in which experimental *Mtb* samples were exposed to. Accordingly, our interdisciplinary research team was able to verify the dose of radiation each sample received in order to confidently attribute the levels of specific biomarkers to a definite dose of radiation.

Specifically, a comprehensive dosimetry analysis of the sample chamber of a JL Shepherd Model 81-14 sealed source ^137^Cs irradiator was performed. The irradiator was initially commissioned in November of 1976 with an original source activity of 6000 Curies. The schematics for the irradiator design were limited but allowed for a rudimentary approximation of the geometrical attributes of the irradiator. Neglecting collimation and scattering effects, that would naturally increase the radiation dose rate within the sample chamber, a line source calculation provided an estimated dose rate of 3.25 Gy·min^-1^. Physical measurements with an ion chamber, thermoluminescent dosimeters, (TLD) and Fricke dosimetry provided dose rates of 7.19 Gy·min-1, 7.56 Gy·min-1, and 5.2 Gy·min-1 (Table 1) respectively. This dosimetry evaluation provided an estimate of the radiation dose rate within the sample chamber, allowing for the exposure duration to be calculated to deliver a defined radiation dose. Specifically, this allowed for radiation dose verification delivered to Mtb samples used to correlate biological effects or subsequent biomarkers to the delivered dose of radiation.

**Table 1.** Average sample chamber dose rates determined by Fricke dosimetry.

| Replicate | *Slope* | $R^2$ |
| --- | --- | --- |
| 1 | 5.18 | 0.99 |
| 2 | 4.96 | 0.965 |
| 3 | 5.49 | 0.981 |

Alamar blue dye contains resazurin, a blue compound which is reduced to a pink fluorescent resorufin when the oxygen substrate is used for cellular respiration as a precursor for cellular metabolic activity (Supplementary Figure 1). Since resazurin has a relatively high reduction potential of +380mV at pH=7, 25°C, it easily accepts electrons during cellular respiration, from several metabolic cofactors such as NADH and FADH^35,36^. This dye is useful for a variety of applications, such as microbial detection assays in food contamination QC, and popularly as a measure of cell viability for cells treated with test compounds^35^. This assay has been particularly popular in measuring the viability of *Mycobacteria* following treatment with a wide range of compounds^35^. However, it has been recently demonstrated that for *Mtb* the Alamar blue assays have a wider range of sensitivity, especially for accurate drug susceptibility testing as compared to other viability assays such as nitrate reductase assays, and MGIT 960^37^. This may more broadly imply, that there exists a higher probability of misleading results from alamar blue assays. For example, positive blue results may be due to contamination of media.

It was necessary to correlate these markers of cell death to dose rate as determined by our multitude of traditional dosimetry techniques. Biological responses to lower doses of radiation provide valuable insight into how radiation damage specifically inactivate certain pathogens, and D_10_-values offer a reliable standard for similar organisms. D_10_-values from dose-response modeling of inactivation offers a means for comparing the relative success of various inactivation methods^26,38^. Dose-response modeling is important for describing pathogen susceptibility to ionizing radiation exposures^26,38-41^, with methods for generating D_10_ models dating back nearly 50 years^41^.

We generated a dose-response curve based on increasing radiation doses delivered to *Mtb* standards, and the consequent decreases in colony forming units (CFU) /ml and cellular respiration. The subsequent dose response curve provides a generalized biological method to correlate the purported radiation dose a sample of *Mtb* receives, to select viability markers. Accordingly, the comprehensive dosimetry assessment further specified the approximate dose-rate, which could be associated to the measured reduction of CFU/ml. This study therefore provided a survival curve and D_10_-value for *Mtb* based on selected viability markers of CFU/ml reduction, supported by a multitude of radiation dosimetry techniques. Specifically, an association between radiation doses and the reduction of CFU/ml from a given sample space, and from that survival curve generate a D_10_-value. This work was crucial in correlating the radiation dose to the biological response in terms of CFU/ml, and alamar blue dye reduction, as markers of reduction in cell viability. The reproducibility and accuracy of the curve we generated, implies that the protocol will be effective elsewhere. Theoretically, the methodology in this work can be applied elsewhere, in different facilities, with substantial success and could help prevent the accidental release of respiratory pathogens into an improperly contained environment, which pose a significant risk to community and global public health^42,43^.

## Methods

### Fricke Dosimetry

For each measurement, the Fricke dosimetry solution was prepared, exposed to the source, and analyzed on the same day. Preparation of the Fricke dosimetry solution and washing of the dosimeter vessels followed the ISO/ATSM standard protocol^44^. Samples were placed into the irradiator’s sample chamber as shown in Supplementary Figure 2. Each designated exposure duration was therefore comprised of a five-sample average. An infrared thermometer was used to measure the temperature of the Fricke solution both prior and following radiation exposure while the samples were still held in the irradiator’s sample chamber. Following radiation exposure, the Fricke solution from each sample was analyzed using a Beckman DU 530 UV Spectrometer set to a wavelength of 303 nm. The temperature of the Fricke solution was recorded during spectrometry to account for temperature variance during exposure and analysis according to the ISO/ATSM protocol^44^.

### Ion Chamber Dosimetry

A Fluke 35040 Advanced Therapy Dosimeter was used to measure the dose rate at two positions within the irradiator’s sample chamber. Measurements were taken for an exposure duration of 30, 60, and 180 seconds at each position with biases of -300, 150, and 300 Volts.

### Thermoluminescent (TLD) Dosimetry

TLD-100 chips, initially calibrated on site with a NIST traceable ^137^Cs source, were annealed for 30 minutes at 360°C before being placed in the same locations as the Fricke dosimeters within the sample chamber. The TLD chips were exposed to the radiation source for 0.1, 0.2, and 0.3 minutes. Following radiation exposure, the TLD’s were analyzed with a Model 3500 Manual TLD Reader with a time temperature profile consisting of 100°C preheat temperature, slope of 10°C•s^-1^, hold time of 26.6 s, and a max temperature of 300°C.

### Sample Preparation for Mtb Dose Calibration

Working stocks of *Mtb* were generated by culture on 7H11 media incubated at 37°C. After four weeks of growth, plates were transferred to glycerol-alanine-salts (GAS) media + 20% glycerol, and stirred on a magnetic stir plate for another 4-6 days. 1 ml aliquots were transferred to 1ml cryovials and stored at -80°C until use. Ninety working stocks were thawed and distributed: three stocks each into 30 5×5 fibreboard cryogenic freezing boxes (VWR) in three positions (top left, true center, and bottom right). The remaining 22 positions not containing cells, were filled with 1.2 ml cryovials containing 1 ml of GAS media + 20% glycerol. Four sample replicates were prepared and used for each selected radiation dose. (n=4 per dose) Additionally, for each treatment group a control box was prepared to be placed proximal to the irradiator. Control boxes were placed in a shielded location at the irradiation facility in order to establish a control for packaging and environmental conditions that may impact bacterial growth. The four remaining vials of *Mtb* were kept at -80°C, to be used as positive controls in CFU and Alamar Blue experiments.

### Mtb Exposure to Ionizing Radiation

Sample boxes were wrapped within three autoclave bags, to ensure triple-containment of potentially viable pathogenic organisms. For the duration of each radiation exposure, similarly prepared control samples were placed within the irradiation facility proximal to the irradiator, but were shielded from the elevated radiation field produced by the unshielded ^137^Cs source. During radiation exposure, sample boxes were placed within the sample chamber of the irradiator. According to the results of the radiation dosimetry assessment, the prepared *Mtb* samples were treated with radiation doses of 0.001 Mrad, 0.005 Mrad, 0.010 Mrad, 0.050 Mrad, 0.100 Mrad, or 0.150 Mrad. For this study, each trial dose was replicated four times. Following radiation exposure, samples were transported back to the BSL-3 laboratory, and frozen at -80°C. Although loss from an additional freeze-thaw was anticipated, a post-treatment freeze-thaw is consistent with the current on site protocol for *Mtb* inactivation.

### CFU Plating and Counts

Boxes were thawed, and the contents of each sample vial were transferred to individual 1.7 ml microcentrifuge tubes. Each tube was spun on a microcentrifuge at 3.75K for 10 minutes, and the supernatant was removed after centrifugation. 1 ml of phosphate-buffered saline (PBS) was added to each tube, followed by an additional centrifugation at 3.75K for 10 minutes. PBS supernatant was removed, and cells were resuspended in 1 ml of 7H9 media with OADC and 0.05% Tween 80. Each tube was placed in an ultra-sonic water bath and sonicated 30 seconds on, and 30 seconds off, three times. In a 24-well sterile plate, each of the sonicated samples were diluted in a 1:10 series to the 10^−6^ dilution. Dilutions were made with 100 µl of sample in 900 µl of 7H9 media with OADC and 0.05% Tween 80 and plated in triplicate on 7H11 agar in quad petri dishes. As part of the colony formation assay, plated samples were incubated at 37°C for 16 days prior to quantifying colony formation. The remaining 900 µl from each sonicated sample, were transferred to their own sterile glass tube containing a sterile stir bar. Samples were placed on a stir plate for 72 hours at 37°C in preparation for Alamar blue viability testing.

### Alamar Blue Viability Testing

Samples were removed from incubation and 200 µl of each was added in triplicate to a 96-well plate. Samples were read at optical density wavelength of 600nm (OD_600_) to standardize concentration. Samples were then diluted to standardize to an OD_600_ of 0.1, and serial diluted in 7H9 + OADC + 0.5% Tween media 1:10 out to an OD_600_ of 0.001. 20µl of Alamar Blue dye (Invitrogen) was added to each sample well. Initial absorbance measurements were performed at dual wavelengths of 570nm and 600nm, representing the 0hr timepoint. Plates were wrapped in protected from light, incubated at 37°C, and absorbance measurements were repeated at 24h and 48h timepoints. A fitted linear regressions model was applied to each series of measurements, and the corresponding slope was used for comparative analysis. Regressions were applied regardless of R^2^-values. Slopes of each sample regression were compared to the respective facility control sample using non-parametric Mann-Whitney t-tests from GraphPad Prism 7.04.

## Results

### Summary of Dosimetry Results

The Fricke dosimetry results shown in Table 1 were obtained from three trials. Each trial consisted of 4 exposure durations, with 5 dosimeters for each exposure. Each dosimeter consisted of 5 ml of Fricke solution held in a glass liquid scintillation vial resulting in a fill height of 1.1 cm. From these measurements, an average dose rate of 5.20 Gy·min^-1^ was determined.

Two positions in the chamber were considered and it was found that Position 1 had an average dose rate of 6.73 Gy·min^-1^, and Position 2 had an average dose rate of 7.87 Gy·min^-1^. Once the initial ion chamber measurements were corrected for the radiation dose associated with the transition of the source, an average dose rate of the sample chamber was approximated to be 7.19 Gy·min^-1^ according to the results of the ion chamber.

TLD-100 chips, placed in the same locations in which Fricke dosimetry was preformed, and were exposed to 3 different durations of radiation exposure. For each exposure duration, measurements were repeated 3 times, providing an average dose rate at the bottom of the sample chamber of 7.56 Gy·min^-1^.

### Linear Regression and D_10_ Value from CFU/ml

A linear regression model was fitted to the treatment doses affect on CFU/ml with an R^2^-value of 0.9941. The dose-response curve, generated as a linear regression on a log-scale ordinate, predicts a D_10_ value of 0.029 Mrad or a radiation dose of 0.029 Mrad necessary for a one log, or 90% reduction in CFU/ml (Figure 1).

**Figure 1.**
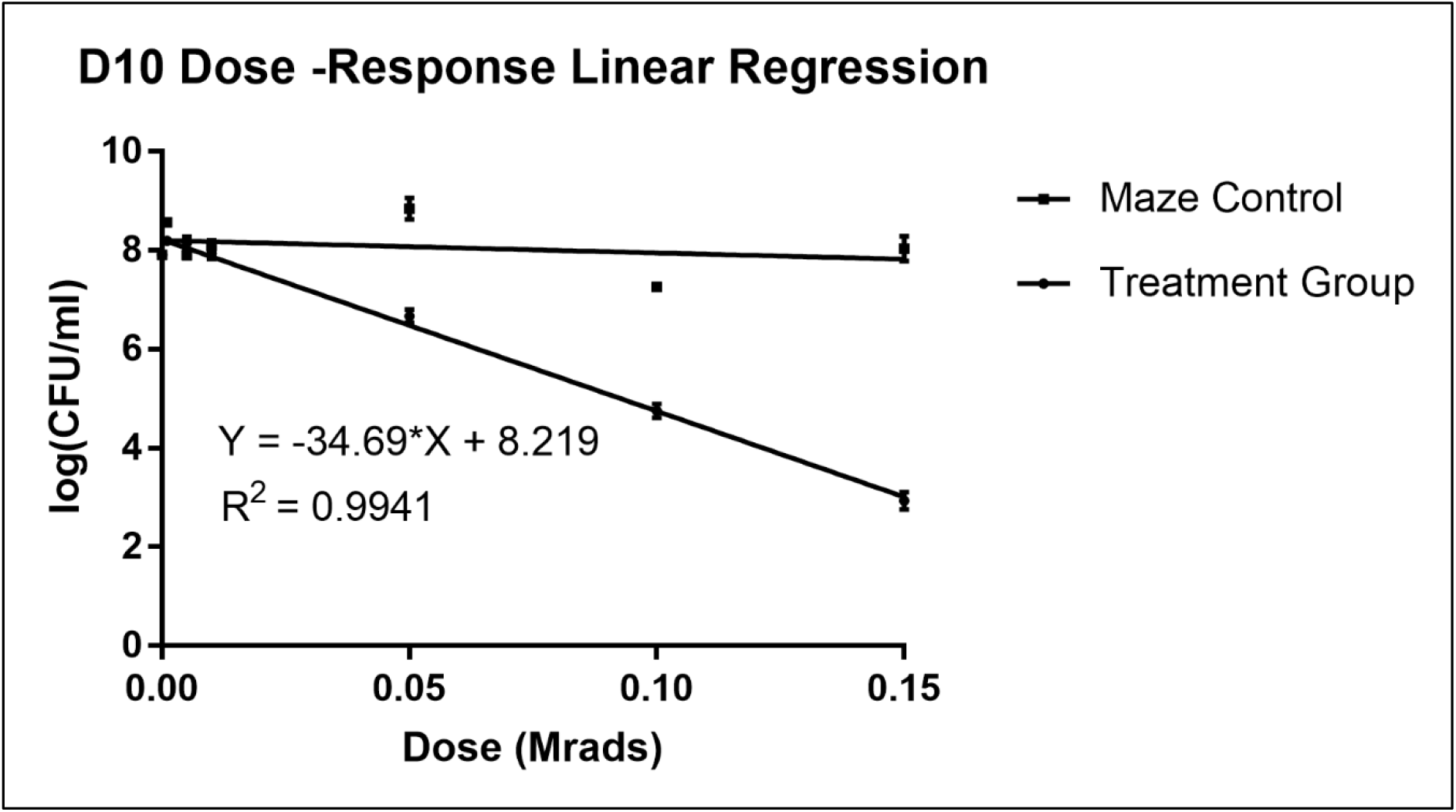
Viability of *Mtb* exposed to ionizing radiation. Mtb H37Rv was exposed to increasing doses of ionizing radiation and analyzed for CFUs on 7H11 agar. Dose Response generated based on CFU/ml after treatment, with slope and R2 of Treatment Group given. Slope indicates D10 value of 0.029 Mrad, that being the dose which leads to 1 log, or 90% reduction in CFU/ml. Regression slope test indicates with a p-value < 0.0001 that the difference in slopes is not equal to zero, indicating a significant difference between slopes.* indicates p-value<0.05.

Reduction in CFU/ml was measured according to the box position parameters described within the methods. No significant difference between CFU/ml reduction and vial position were found, implying even distribution of irradiation (Figure 2).

**Figure 2.**
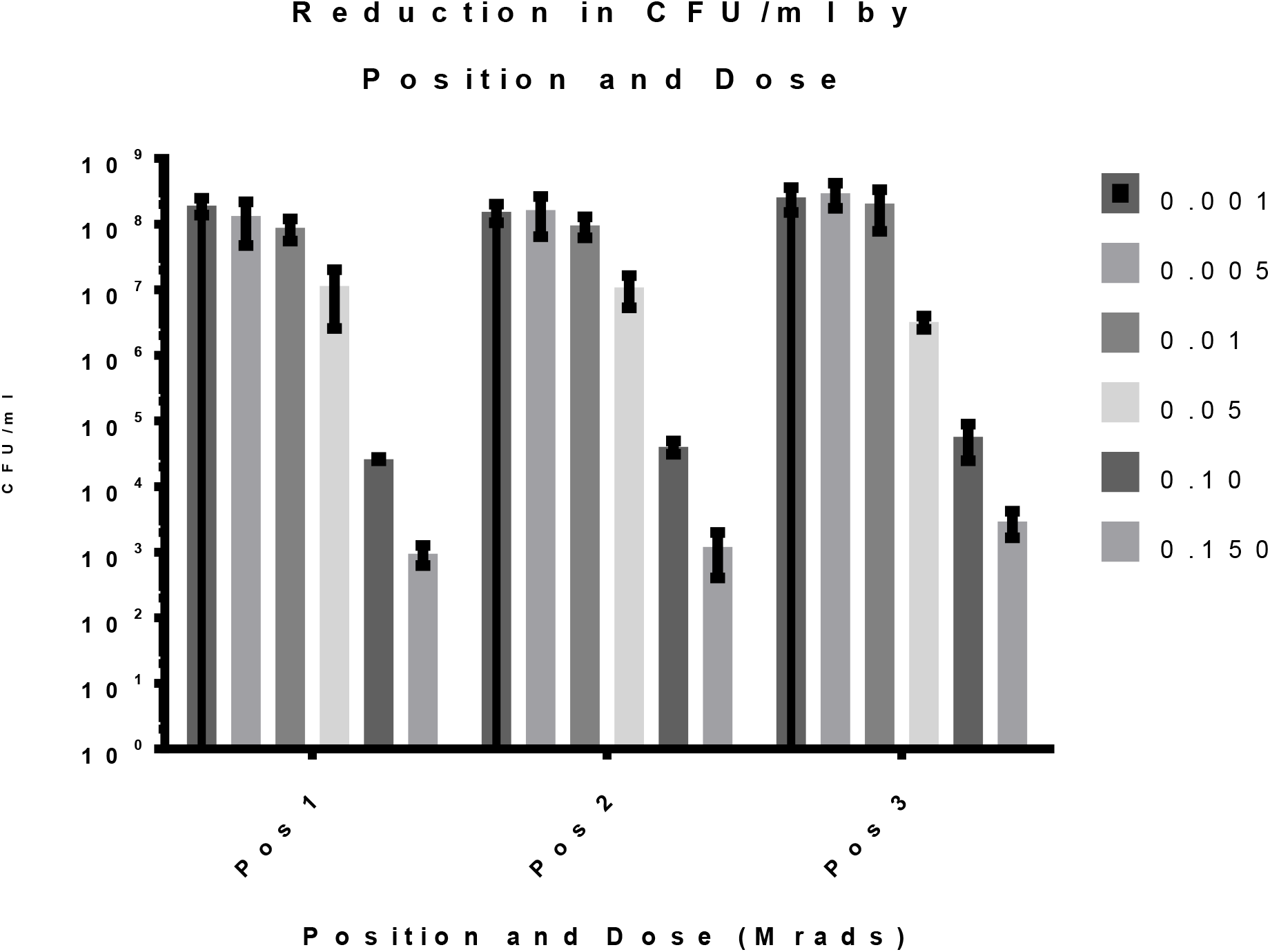
Position-Based Reduction of CFU/ml. CFU/ml measurements were collected from three different positions with each sample box. Reduction of CFU at each positions relative to increased radiation dose was tested. No statistically significant difference in reduction at any timepoint between positions was observed.

### Alamar Blue

General enzymatic activity and organism viability were compared by according to the slop of the linear regression model fit to the Alamar Blue results. Samples exposed to higher doses of radiation yielded reduced accumulation of Alamar Blue dye. Shown in Table 2, a significant variance of Alamar Blue dye reduction was not observed until samples were exposed to a radiation dose of 0.15 Mrad (Table 2). This dose of radiation correlates with a 5.2 log reduction in CFU/ml shown in Figure 1. A significant difference, was found between the irradiation facility control samples and the samples that received a 0.005 Mrad dose of radiation.

**Table 2.**
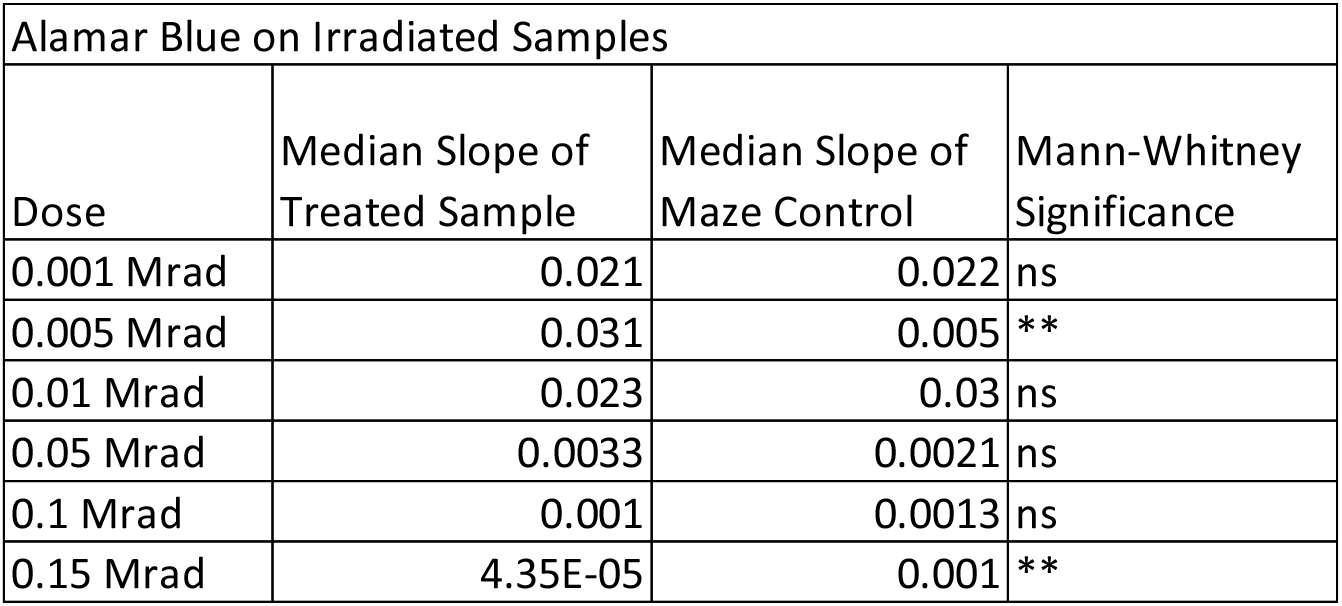
Alamar Blue Slopes of Reduction. Alamar Blue dye assays were performed at 0h, 24h, and 48h timepoints, and slopes were applied to changes in absorbance as measured by 570nm over 600nm. Mann-Whitney nonparametric t-tests were used to compare median slopes generated in Alamar Blue dye assays, between samples exposed to ionizing radiation and irradiation facility control samples. ** = p < 0.005

## Discussion

### Dose Rate Estimate from a Simplified Model

The detailed schematics, or operational mechanism, of the JL Shepherd Model 81-14 sealed source ^137^Cs irradiator were not available at the time of dosimetry. Accordingly, some approximations needed to be applied. The initial dose rate that was caluclated using a line source equasion neglected the radiation dose associated with the backscatter of collimation of the radiation; therefore it was expected that this calculated dose rate would greatly undestimate the true dose rate within the sample chamber of the irradiaor. Nevertheless, this initial caluation served as an approximation of the dose rate in order to estimate the time required to deliver threshold radiation doses for subsiquent dosimetry measuments.

The sample chamber of the irradiator was measured to have a 16 cm width, 9 cm length, and 6.5 cm depth. During operation of the irradiator, the source transitions from a shileded to unshilded position above the sample chamber. Therfore, the dose rate at the top of the sample chamber, which is closer to the source, would have a higher dose rate as compated to the bottom if the sample chamber. For this study, the desired application of the irradiator was for inactivation purposes, therefore dosimetry was performed at the bottom of the chamber to provide the most conservative estimate of the lowest dose rate a sample would receive during radiation exposure.

For each applied dosimetry method, multiple locations on the bottom of the sample chamber were analysed in ensure an approximately homogeneous radiation field. Fricke dosimetry, and radiation dose mearuments with TLD’s were performed at each corrner of the sample chamber, along with the very center sample chamber. Measurments with the ion chamber were taken either on the left or right side, termed position 1 or position 2, of the sample chamber. From the dosimetry results, the higest dose rate was found to be in the center of the sample chamber, position 2, of the sample chamber had a slighetly higher dose rate as compared to position 1. Shown in Figure 2, the minor dose rate variences determined within the sample chamber were not significant enough to alter the biological response of the *Mtb* samples exposed to ionizing radiation. Accordingly, the average dose rate determined for the sample chamber was deemed sufficient to support the conclusions of the dose response assay.

Variability between dosimetry methods was to be expected. Fricke dosimetry is considered a dosimetric standard, but the results are easily influenced by external factors and impurities. To identify potential errors, fresh batches of the Fricke solution were prepared for each trial utilizing different sources of sulfuric acid and ferrous ammonium sulfate. Due to the high activity of the source, and corresponding high dose rate within the sample chamber, only short exposure durations were measured with the ion chamber and TLDs. Initial measurements with the ion chamber included the radiation dose associate with the transition of the source for the shielded to unshielded position. Although the radiation dose associated with the transition of the source was found to be minimal, the initial experimental design in which short radiation exposures were used to determine the dose rate, had the potential to misrepresent the dose rate to be greater than the actual dose rate if the radiation dose of the source transition was not accounted for. Therefore, it is recommended that when performing radiation dosimetry an understanding of the working mechanisms specific to the radiation device is necessary to not misrepresent the dosimetry results. This would be especially pertinent due to significant safety implications for radiation devices used for pathogen inactivation. Additionally, due to the variation between each of the results of the applied dosimetry methods, a comprehensive dosimetry assessment should not rely on the results of a particular methodology, rather a combination of dosimetry methods provides a more comprehensive estimate of the actual dose rate. More so, the application of the radiation device must also be taken into account.

### Mtb Dose-Response and D_10_ Value

In this study, we employed methods for calibration of irradiation devices used to inactivate pathogens. Not only were the data points collected highly reproducible, but the applied linear regression model accounts for nearly all statistical variance, thus indicating that the model is sutible for future experiments. This D_10_ method and model can be used to describe the effects of ionizing radiation to organisms treated at determined doses.

A radiation dose of 0.050 Mrads was shown to reduce cell culturing viability by at least 90%. The model indicates that a radiation dose of 0.029 Mrad would be sufficient for a one log or 90% reduction in CFU/ml. For future calibration efforts, comparisons in CFU/ml reduction can be made on the basis that a radiation dose of 0.029 Mrads generates a one log reduction in CFU/ml (Figure 1). Practically speaking, this means that we have a robust D_10_ value for future reference, as well as a methods for conduction this kind of experiment for future applications. A reliable D_10_-value for *Mtb* treated at determined doses, will also allow for simple calculation of the effect of treatments under varying conditions, to cell death and loss of generalized enzymatic activity.

Irradiation position data (Figure 2) was compelling in its demonstration that position on the irradiation device does not influence the reduction of CFU/ml that we observed. Each dosimetry method showed dose rate variances within the sample chamber, but that was not observed by the biological samples. This indicated that the using CFU/ml is not as sensitive of a method for radiation dosimetry. Therefore, radiation dosimetry must be determined independently of the treatment of a biological sample to determine potential variances within a given sample chamber.

Alamar Blue results showed a threshold radiation dose of 0.15 Mrads in order to elicit a statistically significant reduction of dye. This correlates to an approximate 5.2 log reduction in CFU/ml as modeled in Figure 1. Previous literature has long demonstrated that a >6 log reduction in CFU/ml is necessary for confidence for the inactivation of viable pathogens^4^. The Alamar Blue dye assay is consistent with this standard, and was shown to be more sensitive as compared to CFU/ml by only demonstrating a significant difference once the CFU reduction neared a 6 log decrease. In other words, a higher radiation dose and consequent reduction in viable cells to observe differences between treatment and control groups in Alamar Blue assays, compared to CFU assays. Future work can be done to determine exact expected reductions in Alamar Blue colorimetric changes associated with a >6 log reduction. The difference in viability assay results may be indicative of cells which are no longer culturable but remain metabolically active specifically when exposed to low doses of radiation. This type of cell-state can be described as viable but non-culturable (VBNC), and it has significant implications for low-dose inactivation studies. VBNCs induced by extracellular stress are described in *Mycobacterium spp*.^45,46^ as well many other bacterial species^47,48^. They are characterized as cells which have lost their ability to grow in media, while maintaining structural integrity, some metabolic function, cell wall integrity, protein function, and so on^45-48^. Cells in such a state would appear inactive in cultivation-based viability assays, due to the loss of culturability expected from live laboratory-strain cells. However VBNCs in dormant, or even pre-death states may still generate responses in metabolic-based viability assays. More concerning, VBNCs are associated with reactivation and maintained virulence in pathogenic organisms.

Interestingly, a significant difference between the samples exposed to a radiation dose of Mrads and the control groups was observed. We hypothesize that this was most likely due to an increase in metabolic activity, as opposed to the decreases in activity observed in consequent higher doses. This could be due to the dose inducing stress conditions, particularly oxidative stress, upregulating DNA-repair mechanisms which have evolved to specifically react to distinct types of damage^49^. Additionally, *Mycobacteria* have been demonstrated to be uniquely adept at surviving a wide range of threats to cellular integrity and survival^50^. Often a number of heat shock proteins such as ACR and HSP-3, as well as KatG stress-related genes, whiB-like genes and σ-factors are attributed to this survival characteristic^50,51^. The result of initiation of these cascades at low levels may result in enhanced growth and metabolism. Additionally, increases in transcriptomic and metabolic activity are associated with oxidative stress at low and intermediate levels, with no negative effect on growth^52^. Our findings at the 0.005 Mrad dose are consistent with these previous observations and *Mtb* exposed to low radiation doses could actually enhance the growth and metabolic rate of the organism. Low-level radiation would therefore cause an hormetic effect on *Mtb*. Hormesis induction by low concentrations of nitric oxide has been previously described in *Mtb*^53^, indicating that the response is not unprecidented. However, similar research has not been conducted using low-doses of ionizing radiation. One important future direction of this work will be to interrogate this possibility. Use of thawed samples (as we did), or the likely conditions where samples thaw on instrument, may impact dosing as well. Metabolomic and proteomic studies using thawed sample, frozen sample, or sample on a cooling chamber, will need to be conducted in the future to further demonstrate the effect of low doses on *Mtb* and other pathogenic cells.

Our work has led to a reproducible method and standard for comparative studies to study the effect of ionizing on cell viability. This work has not addressed other issues such as the safety and cost associated to utilizing radiation, and specifically a sealed source irradiator, for the inactivation of pathogens, For that reason, future studies may focus on comparing radiation using the D_10_ method to alternative inactivation techniques in order to evaluate robust inactivation strategies that are safer and more cost-effective than gamma-irradiation. It will be paramount to further evaluate such methods as well, for their maintenance of structural and antigenic attributes of inactivated pathogens.

## Concluding Remarks

Inactivation techniques, despite their previously demonstrated successes or long-standing reputations, should always continue to be interrogated and improved upon to better understand the negative attributes associated with each modality. Inactivation of pathogens from large-scale culture plays a crucial role in research, food safety, and transfusion of plasma products. However, the consequences of incomplete inactivation can be devastating, therefore improving methodologies used for the inactivation of a pathogen, and updating corresponding protocols will help improve safe manipulation of pathogens decreasing the likelihood of laboratory acquired exposures or infections.

The results obtained by different dosimetry methods in the course of this study are consistent for each measurement method, but display a marked variation between the different methods. It is important to recognize that this variation might be indicative of the inherent uncertainty for each measurement modality. While the simple mean of the results obtained by the three different measurement protocols would indicate a dose rate of 6.6 Gy·min^-1^, the associated uncertainty of 1.3 Gy·min^-1^ shows that any particular value for the dose rate used to assess the total dose applied needs to be evaluated explicitly recognizing these uncertainties. It is important to understand that a true value for the dose rate could lie anywhere within the range of approximately 5 Gy·min^-1^ to approximately 8 Gy·min^-1^. We conclude that we must ensure consistency of the radiation field for future applications, rather than emphasize one particular value for the dose rate.

Regardless of how dose rates are determined, they must also be associated with reduction in viable cells if radiation is to be used for complete inactivation of cells. The application of a radiological dose to a given pathogens is ambiguous, in the context of inactivation research, without a means to associate dose to biological markers of death. A kill-curve and subsequent D_10_-value help bridge this gap, in that dose rates can be associated with loss in live cells.

Finally, the potential observation of hormesis induced by low-levels of radiation should be further interrogated. While the phenomenon has been described in *Mtb* previously, it has not been investigated using ionizing radiation. This represents a plausible, and novel hypothesis for radiation-induced hormesis in *Mtb*.

## Acknowledgements

We would like to acknowledge Mike Weil for his helpful contributions to this study, as well as Anne Simpson for technical assistance. Funds for this study were provided to AB and KMD through Colorado State University.

